# Multi-omics investigation of tacrolimus nephrotoxicity

**DOI:** 10.1101/2021.07.29.454229

**Authors:** Hassan Aouad, Quentin Faucher, François-Ludovic Sauvage, Emilie Pinault, Claire-Cécile Barrot, Hélène Arnion, Pierre Marquet, Marie Essig

**Author notes:** **Correspondance :** Pierre Marquet, Service de Pharmacologie, Toxicologie et Pharmacovigilance, Centre Hospitalier Universitaire, 2 rue Martin Luther King, 87042 Limoges, France. These authors have contributed equally to this work and share last authorship (Pierre Marquet, Marie Essig). Primary laboratory of origin: U1248 IPPRITT, Université de Limoges, INSERM, Limoges, France.

## Abstract

**Background:** Tacrolimus, an immunosuppressive drug prescribed to a majority of transplanted patients is nephrotoxic, through still unclear mechanisms. This study aims to evaluate the impact of tacrolimus on a lineage of proximal tubular cells using a multi-omics approach.

**Methods:** LLC-PK1 cells were exposed to 5 M of tacrolimus for 24h. Intracellular proteins and metabolites, and extracellular metabolites were extracted and analysed by LC-MS/MS. The transcriptional expression of the dysregulated proteins PCK-1, FBP1 and FBP2 was measured using RT-qPCR.

**Results:** In our cell model, tacrolimus impacted different metabolic pathways including those of arginine (e.g., citrulline, ornithine) (p < 0.0001), amino acids (e.g., valine, isoleucine, aspartic acid) (p < 0.0001) and pyrimidine (p<0.01). In addition, it induced oxidative stress (p < 0.01) as shown by a decrease in total cell glutathione quantity. It impacted cell energy through an increase in Krebs cycle intermediates (e.g., citrate, aconitate, fumarate) (p < 0.01) and down-regulation of PCK-1 (p < 0.05) and FPB1 (p < 0.01), which are key enzymes in gluconeogenesis. Apart from glucose synthesis, gluconeogenesis is an important process in kidney mediated acid-base balance control.

**Conclusion:** The variations found using this multi-omics approach clearly point towards a dysregulation of energy production in epithelial cells of the renal tubule, and potentially of their functions, that may be implicated in tacrolimus nephrotoxicity in the clinics.

## Introduction

Tacrolimus is an immunosuppressive drug widely used to prevent graft rejection after solid organ transplantation and to treat autoimmune diseases (Brunet et al., 2019). Its whole blood concentrations should be within a narrow therapeutic range of 4-15 ng/ml to avoid underexposure and increased risks of rejection, and overexposure known to entail acute and chronic nephrotoxicity. Acute nephrotoxicity, linked to tacrolimus blood level >20 ng/ml, is caused by hemodynamic perturbations and is reversible (Böttiger et al., 1999). In contrast, chronic nephrotoxicity is an irreversible decline of renal function that can appear along time even in patients exposed to (low) pharmacological levels of tacrolimus.

The mechanism of tacrolimus nephrotoxicity is still not fully understood. Many studies were conducted on proximal tubular cells and linked nephrotoxicity either to a mitochondrial toxicity characterized by functional and structural perturbation of the mitochondria (Lim et al., 2019a; Yu et al., 2019), or to increased oxidative stress caused by an increase of intracellular H_2_O_2_ levels or a decrease in MnSOD, an antioxidant enzyme (Lim et al., 2017; Zhou et al., 2004). Autophagy was also found to be disrupted in cells exposed to tacrolimus, with an accumulation of autophagy vesicles in the cytoplasm (Lim et al., 2019b; Zheng et al., 2020). Furthermore, cells exposed to tacrolimus were also reported to lose some transporter-related functions (Secker et al., 2019).

In this multi-omics study, we explored the intracellular and extracellular modifications of LLC-PK1 cells incubated with tacrolimus, in order to decipher the pathways involved in its toxicity on the proximal tubule. We confirmed that tacrolimus induces an oxidative stress and found that it also affects the arginine and energetic metabolisms, as suggested by increased quantities of intermediates of the citric acid cycle and down-regulation of PCK-1 and FBP1, which are gluconeogenesis-limiting enzymes. In addition, tacrolimus modified the metabolism of purine bases.

## Materials and methods

### Chemicals and reagents

Dulbecco’s Modified Eagle’s Medium (DMEM)-Ham’s F12 (1:1, 31331), Fetal Bovine Serum (10500), 1 M HEPES (15630), 7.5 % Sodium bicarbonate (25080), 10,000 UI/mL Penicillin/ Streptomycin (15140), Dulbecco’s Phosphate Buffer Saline (14190), Superscript II RT (180064022, Invitrogen™) and BCA protein assay KIT (23225, Pierce™) were purchased from ThermoFisher Scientific (Illkrich-Graffenstaden, France). Sodium selenite (S5261), insulin (I4011), triiodothyronine (T6397), dexamethasone (D4902), human apo-transferrin (T1147), desmopressin (V1005), tacrolimus (F4679), 2-isopropylmalic acid (333115), DTT (1.4-Dithiothritol) (D0632), Urea (U5378), Iodoacetamide (I1149), AmiconUltra-0.5 centrifugal filters (UFC5010) and DNase I Kits (AMPD1) were purchased from Sigma-Aldrich (St. Quentin Fallavier, France). NuleoSpin® RNA/Protein extraction kits (740933.50) were purchased from Macherey-Nagel (Hoerdt, France). Sequencing grade modified trypsin (V5111), RNasin® ribonuclease inhibitor (N2511), random primers (C118A) and MTS (G3581) were obtained from Promega (Courtaboeuf, France). QuantiFast® SYBER® Green PCR Kits (204054) were purchased from Qiagen. HLB oasis 3cc 60mg cartridges (WAT094226) were obtained from Water (saint Quentin en Yvelines, France).

### Cell culture conditions

LLC-PK1 (Lilly Laboratories Porcine Kidney-1) porcine proximal tubule cells (ATCC-CL-101, ATCC, Manassas, VA) were expanded in 75 cm^2^ flasks at 37 °C with 5 % CO_2_ and passed once confluence was reached. The culture medium consisted in a 1:1 DMEM-Ham’s F12 mix supplemented with 5 % FBS, 15 mM HEPES, 0.1 % Sodium bicarbonate, 100 UI/mL Penicillin / Streptomycin and 50 nM Sodium selenite. LLC-PK1 cells were cultured between passage 7 and passage 20.

LLC-PK1 were seeded in 6-well plates and expanded up to sub-confluence in the routine cell culture medium. Seeded LLC-PK1 sustained serum starvation and were fed with hormonally defined (25 μg/mL insulin, 11 μg/mL transferrin, 50 nM triiodothyronine, 0.1 μM dexamethasone, 0.1 μg/mL desmopressin) fresh medium to engage epithelial differentiation, for 24 hours. After differentiation, two treatment conditions were applied for 24h as follows: i) ethanol 0.5% (control); or ii) tacrolimus 5 μM in 0.5% ethanol. For intracellular and extracellular metabolomics investigations, 3 and 4 independent experiments were performed in triplicate, respectively. For the proteomics study, 6 independent experiments were performed in singlicate.

### Viability test

Cells were seeded in 96-well plates at 25,000 cells/well for 24h, then differentiated for 24h and treated with either 0.5% ethanol (control) or 5μM tacrolimus. Culture medium, with and without tacrolimus, was changed every 24h. Viability was assessed using the MTS test according to the manufacturer’s instruction. Measurement of the absorbance was performed on PerkinElmer EnSpire® Multimode Plate Reader.

Viability was also assessed using flow cytometry. Briefly, cells were cultured in 6-well plates at 500,000 cells/well. After treatment, as described above, cells were washed twice with PBS and detached with trypsin EDTA. Detached cells were then washed twice with PBS and stained with annexin 5/7AAD. Reading was performed on a BD LSRFortessa™ flow cytometer.

### Metabolomics study

#### Sample preparation

Extraction of intracellular metabolites was based on a previously published method (Yuan et al., 2012). Treated and control cells were washed twice with ice cold PBS and then lysed using 3 ml of a mixture of methanol/water 80%/20% volume spiked with 2-isopropylmalic acid at a final concentration of 500 nM (internal standard) and incubated for 20 minutes at -80°C. Every well was then scrapped using cell scrapper, cell lysates transferred to 5 ml Eppendorf tubes and centrifuged at 20,000 g for 5 min at 4°C. One milliliter of each supernatant was then transferred into a 1.5 ml Eppendorf tube and evaporated to dryness in a vacuum concentrator. The extract was then solubilized with 50 μL of MiliQ water and transferred to a vial for mass spectrometry analysis.

Extracellular metabolites were extracted following manufacturer’s instructions (Shimadzu). The culture medium of treated cells was collected and centrifuged at 3,000 g for 1 minute at room temperature to eliminate cell debris. Then, to 100 μL of the supernatant were added 200 μL of acetonitrile and 20 μL of internal standard (2-isopropylmalic acid [500μM]). After homogenization, samples were centrifuged at room temperature for 15 min at 15,000 g. The supernatant therefore was then diluted 1/10 in ultrapure water and transferred into an injection vial for mass spectrometry analysis.

#### Metabolomics LC-MS/MS analysis

Three μL of the suspended extracts were injected into the analytical system. Mass spectrometry analyses were performed using a LCMS-8060 (Shimadzu) tandem mass spectrometer and the LC-MS/MS “Method Package for Cell Culture Profiling Ver.2” (Shimadzu). The mass transitions of additional compounds were added after infusing the pure substances in the mass spectrometer. For each transition analyzed, only well-defined chromatographic peaks were considered. The area under the curve of each metabolite was normalized to the area under the curve of the internal standard (2-isopropylmalic acid).

#### Data analysis and statistics

In every experiment, treated samples were normalized by the corresponding control. MetaboAnalyst 5.0 computational platform (www.metaboanalyst.ca/faces/home.xhtml) was used for all statistical analyses. Univariable analysis was performed using the t-test; p-values were corrected for multiple testing using the False Discovery Rate (FDR) method. Multivariate exploration and unsupervised analysis by Principal Component Analyses (PCA) were performed.

### Proteomics study

#### Sample preparation

For intracellular proteomics, treated and control cells were washed twice with ice cold PBS and then extracted with the NucleoSpin^®^ RNA/Protein extraction kit following manufacturer’s instructions. The extracts were then stored at -80°C until analysis.

For intracellular proteomics, protein content was estimated using the BCA protein assay following manufacturer’s instruction. Thereafter, 50 μg of proteins were diluted in q.s. 200 μL of 8 M urea followed by 20 μL of 50 mM DDT and the samples were then incubated at 56°C for 20 min. Next, 20 μL of 100 mM iodoacetamide were added and the samples incubated in the dark for 20 min. The reduced samples were transferred to an Amicon ultra-centrifugal filter and centrifuged at 14,000 *g* for 15 min. After the first centrifugation, the samples were washed twice by 8 M urea and then twice with 25 mM ammonium bicarbonate. Digestion was performed on the filter by adding 10 μL of a solution of 0.1 μg/μL of trypsin and the mixture was incubated for 3h at 37°C. Peptides were recovered by centrifuging the Amicon filter at 14,000 *g* for 15 min, washing it with 100 μL of 1.5 M NaCl, putting it upside down in the tube and finally centrifuging at 1,000 *g* for 2 min. Solid Phase Extraction (SPE) of the peptides was performed using OASIS® HLB cartridges (Waters) preconditioned with 3 mL of methanol and 3 ml of water/formic acid 0.5%. After loading, the diluted sample was first washed with 3 ml of water/formic acid 0.5% and then with 3 ml of water/methanol/formic acid (94.5/5/0.5, v/v/v). The cartridge was dried for 15 min and elution achieved with 3 ml of acetonitrile/water (70/30, v/v). The eluate was evaporated under nitrogen and the dry residue was dissolved in 100 μl of water/acetonitrile/trifluoracetic acid (TFA) (98/2/0.05, v/v/v). The sample was then filtered on a 0.22 μm spin filter (Agilent) and analyzed by mass spectrometry.

#### Proteomics MicroLC-MS/MS analysis

The peptides resulting from protein digestion were analyzed by microLC-MS/MS using a nanoLC 425 liquid chromatography system in the micro-flow mode (Eksigent, Dublin, CA), coupled to a quadrupole-time-of-flight tandem mass spectrometer (TripleTOF 5600+, Sciex, Framingham, MA) operated in the high-sensitivity mode. Reverse-phase LC was performed in a trap-and-elute configuration using a trap column (C18 Pepmap100 cartridge, 300 μm i.d. x 5 mm, 5μm; Thermo Scientific) and a C18 analytical column (ChromXP, 150 × 0.3 mm i.d., 120Å, 3 μm; Sciex) with the following mobile phases: loading solvent (water/acetonitrile/trifluoroacetic acid 98/2/0.05 (v/v)), solvent A (0.1% (v/v) formic acid in water) and solvent B (water/acetonitrile/formic acid 5/95/0.1% (v/v)). All samples were loaded, trapped and desalted using a loading solvent flowrate of 10 μL/min for 5 min. Chromatographic separation was performed at a flow rate of 3 μL/min as follows: initial, 5% B, increased to 25% in 145 min, then to 95% B in 10 min, maintained at 95% for 15 min, and finally, decreased to 5% B for re-equilibration.

One μg of each sample (equivalent protein content) was first subjected to data-dependent acquisition (DDA) to generate the SWATH-MS spectral library. MS and MS/MS data were continuously recorded with up to 30 precursors selected for fragmentation from each MS survey scan. Precursor selection was based upon ion intensity, whether or not the precursor had previously been selected for fragmentation (dynamic exclusion). Ions were fragmented using the rolling collision energy setting. All DDA mass spectrometry files were searched using ProteinPilot software v.5.0.1 (Sciex) and the Paragon algorithm. Data were analyzed using the following parameters: cysteine alkylation with iodoacetamide, digestion by trypsin and no special factors. The search was conducted using UniProt database (June 2018 release) containing non-redundant proteins of *Sus scrofa*. The output of this search was used as the reference spectral library.

For sample analysis, the equivalent of 1 μg protein content was injected in the analytical system and subjected to data-independent acquisition (DIA) using 60 variable swath windows over the 400-1250 *m/z* range. For these experiments, the mass spectrometer was operated in such a way that a 50-ms survey scan (TOF-MS) was acquired and subsequent MS/MS experiments were performed on all precursors using an accumulation time of 120 ms per swath window for a total cycle time of 7.3 s. Parent ions were fragmented using rolling collision energy adjusted to the *m/z* range window. DIA samples were processed using PeakView v.2.1 (Sciex) with SWATH v.2.0 module and the reference spectral library generated above.

Spectral alignment and targeted data extraction were performed using an extraction window of 15 min and the following parameters: protein identity confidence > 99% with a maximum of 10 peptides per protein and 5 fragments per peptide with 10 ppm error tolerance. Shared and modified peptides were excluded.

#### Data analysis and statistics

Statistical analysis of proteomics results was performed using R. First, each batch of treated and control cells was normalized classically (centered and scaled). Second, delta values were computed from normalized data, as follows:

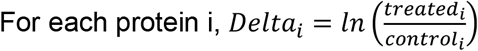

Third, proteins with delta < ln (0.8) or > -ln (0.8), reflecting a decrease or increase of protein expression by more than 20% and identified using more than 2 peptides, were regarded as differentially expressed. Statistical analysis performed using paired t-test with Prism 5.0 software

### RT-qPCR

RNA was extracted together with proteins using the NucleoSpin^®^ RNA/Protein extraction kit. RNA quantification was performed using a Nanodrop® spectrophotometer (ND-1000). To eliminate residual DNA, samples were treated using DNase I Kits (sigma AMPD1). One μg of RNA was then reverse-transcribed into complementary DNA (cDNA) using Superscript II RT, random primers and RNasin® ribonuclease inhibitor (Promega N2511). All these steps were carried out following the manufacturer’s instructions. For qPCR reaction, a mix was prepared for each sample containing 40 ng of cDNA, 2.5μL of a mix of forward and reverse primers (final concentration for each primer: 1μM), 12.5 μL of QuantiFast® SYBER® Green PCR Kit and q.s. 25μL RNAse-free water. The reaction was performed on a Rotor Gene Q (Qiagen) using the following program: 5 min at 95°C followed by 45 cycles of 10s at 95°C, 30s at 60°C. Acquisition was done in green. Fold-change was calculated (2^-ΔΔCt^) and statistical analysis performed using paired t-test with Prism 5.0 software. The complete list of primers used is presented in Supplemental table 1.

## Results

Cell viability was not affected after 24h and 48h of treatment with 5 μM of tacrolimus. At 72h, the MTS test showed a significant decrease of 5 % (p=0.0003) in cell viability, while flow cytometry analysis did not show any difference (Figure 1).

**Figure 1:**
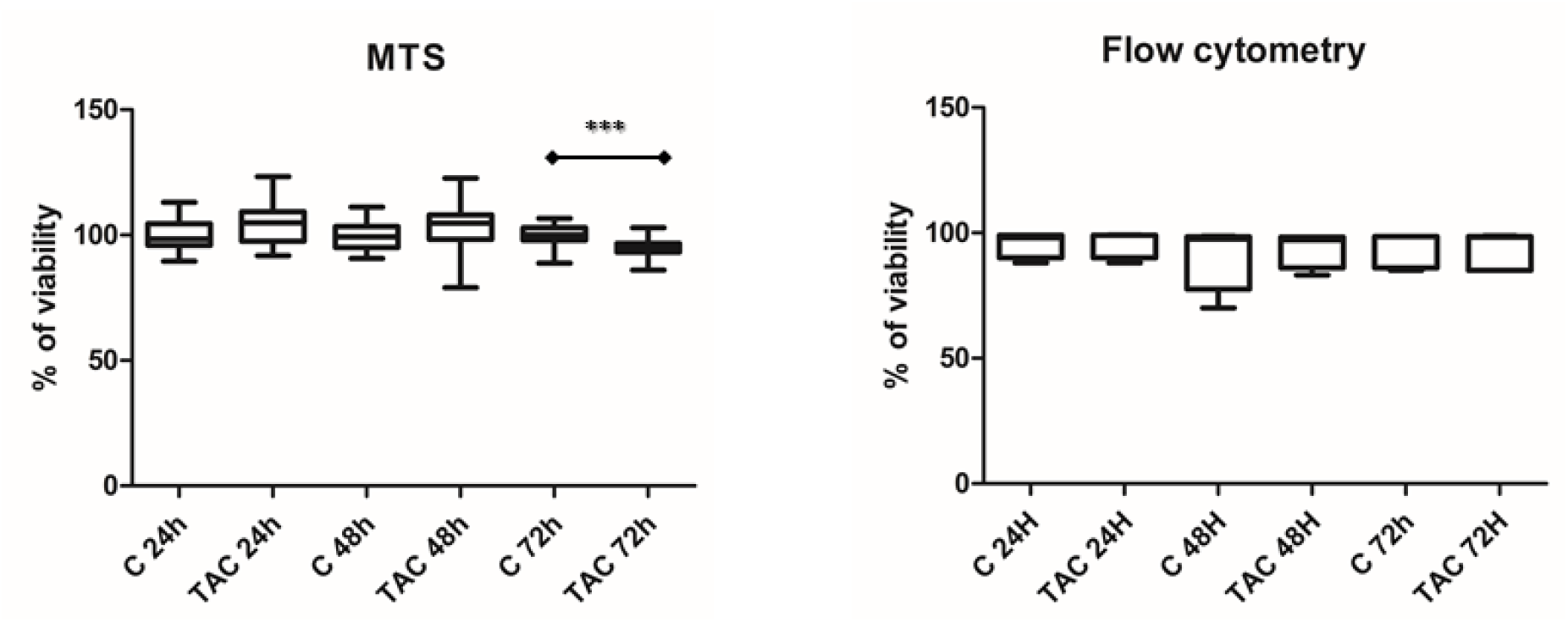
Viability of the tubular proximal cell line exposed to tacrolimus. LLC-PK1 viability after incubation with 0.5% ethanol (control (C)) or 5μM of tacrolimus in 0.5% ethanol (TAC) for 24h, 48h and 72h, assessed using the MTS viability assay (n=18, left graph) and annexin 5/ 7AAD staining (n=5, right graph). Graphs represent the % of viable cells with tacrolimus as compared to control for each incubation duration. *** p < 0.001 by Student t-test.

### Intracellular metabolomics

Seventy-nine metabolites were detected by LC-MS/MS and heat-map analysis clearly discriminated the two conditions (controls or tacrolimus) (Figure 2 and Supplemental table 2). PCA neatly separated the two experimental conditions too, with 47.6% of the variation being explained by the first component and 21.8% by the second (Figure 2).

**Figure 2:**
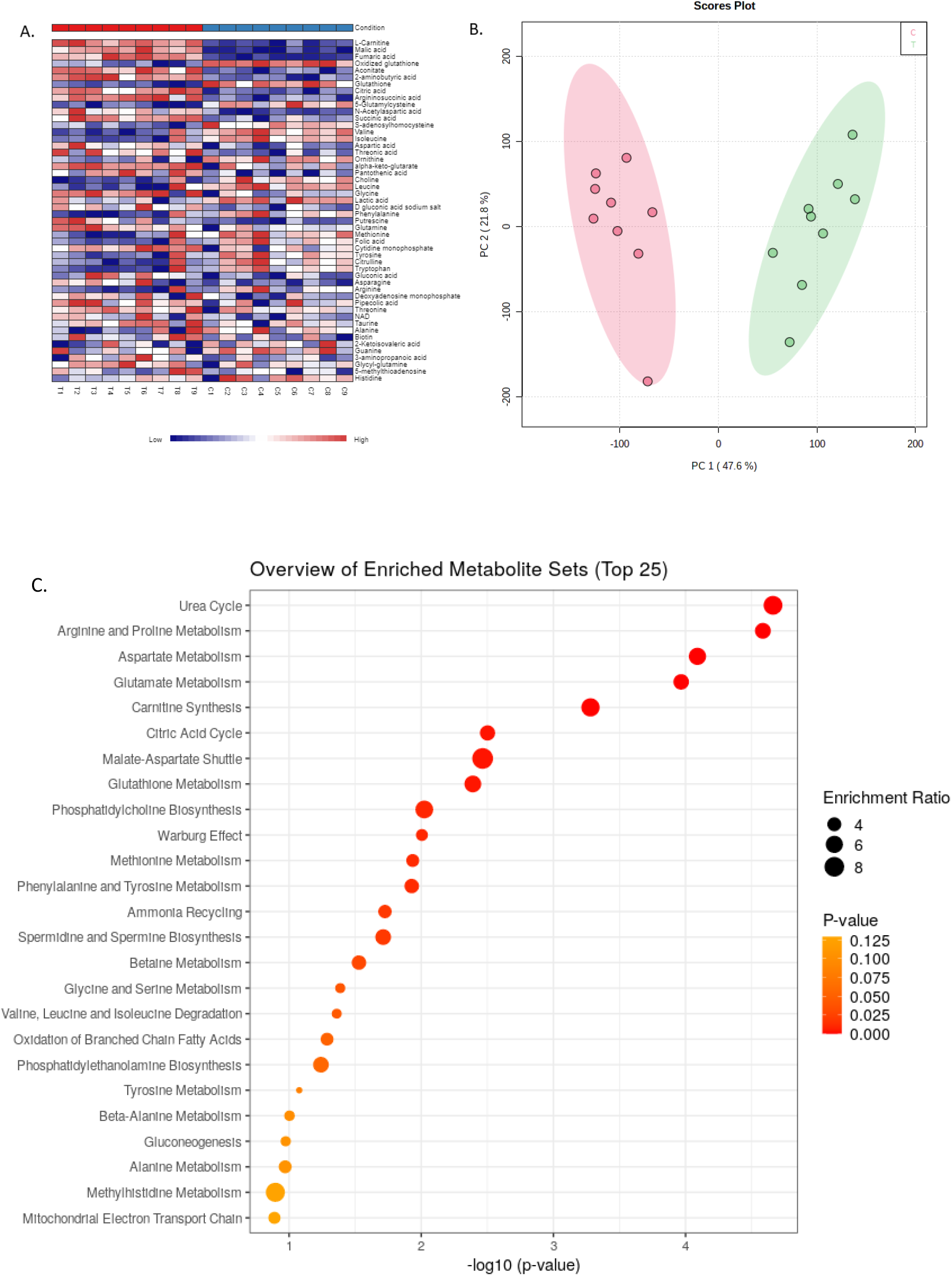
Multivariate exploration of intracellular metabolite variations in response to tacrolimus. Multivariate exploration of intracellular metabolites variations after 24h tacrolimus exposure (n=9). ***A***. Heatmap clustering distinguishing the control condition (C) from tacrolimus-treated cells (T), based on all metabolites detected by LC-MS/MS. **B**. PCA scores plot showing complete separation between groups (C) and (T) with principal components PC1 and PC2 describing 47.6% and 21.8% of the variations, respectively (0 (red circles): 0.5% ethanol (control); 1 (green circles): tacrolimus 5μM). **C**. Dot plot of pathway enrichment analysis based on the intracellular concentrations of metabolites significantly modified in tacrolimus-treated cells as compared to controls (Table 1).

Univariate analysis (t-test) showed that 20 of these metabolites were significantly increased and 15 significantly decreased (p-value<0.05 and FDR<0.05) (Table 1). Most of these metabolites were amino acids, citric acid cycle intermediates, urea cycle intermediates and antioxidant reaction intermediates.

**Table 1.** Fold-change of intracellular metabolites significantly influenced by tacrolimus (n=9) (p-value<0.05)

| Metabolite | Fold-change | P-value (T-test) | FDR |
| --- | --- | --- | --- |
| L-Carnitine | 1.7425 | 8.9464e-11 | 7.0677e-09 |
| Malic acid | 1.4603 | 2.2221e-10 | 8.7772e-09 |
| Fumaric acid | 1.383 | 3.5883e-10 | 9.4491e-09 |
| Aconitate | 1.4654 | 7.0907e-08 | 1.235e-06 |
| 2-aminobutyric acid | 1.2619 | 7.8167e-08 | 1.235e-06 |
| Glutathione | 0.26167 | 1.526e-07 | 2.0093e-06 |
| Citric acid | 1.4012 | 1.8841e-07 | 2.1264e-06 |
| Argininosuccinic acid | 1.4044 | 8.0989e-07 | 7.9976e-06 |
| γ-Glutamylcysteine | 0.38874 | 2.4983e-06 | 2.193e-05 |
| N-Acetylaspartic acid | 1.4012 | 5.9313e-06 | 4.6858e-05 |
| Succinic acid | 1.2442 | 9.4702e-06 | 6.8013e-05 |
| Oxidized glutathione | 0.53446 | 1.206e-05 | 7.9393e-05 |
| S-adenosylhomocysteine | 0.67941 | 2.2869e-05 | 0.00013898 |
| Valine | 0.78981 | 0.0001452 | 0.00081935 |
| Isoleucine | 0.8245 | 0.00018176 | 0.00095725 |
| Aspartic acid | 1.2697 | 0.00044392 | 0.0021918 |
| Threonic acid | 1.3235 | 0.00055436 | 0.0025761 |
| Ornithine | 0.80762 | 0.00067636 | 0.0029685 |
| Oxoglutaric acid | 1.2475 | 0.0008545 | 0.0035529 |
| Pantothenic acid | 1.0773 | 0.00099844 | 0.0037844 |
| Choline | 0.88625 | 0.001006 | 0.0037844 |
| Leucine | 0.85471 | 0.00139 | 0.0049616 |
| Glycine | 1.1806 | 0.0014445 | 0.0049616 |
| Lactic acid | 0.95114 | 0.001715 | 0.0056453 |
| D gluconic acid sodium salt | 1.1862 | 0.001793 | 0.005666 |
| Phenylalanine | 0.89058 | 0.0025434 | 0.0077046 |
| Putrescine | 1.2476 | 0.0026332 | 0.0077046 |
| Glutamine | 1.3081 | 0.0042679 | 0.012042 |
| Methionine | 0.90562 | 0.0048398 | 0.013184 |
| Folic acid | 0.84262 | 0.0055765 | 0.014685 |
| Cytidine monophosphate | 1.1084 | 0.0061877 | 0.015769 |
| Tyrosine | 0.89841 | 0.0077896 | 0.019231 |
| Citrulline | 0.84895 | 0.009764 | 0.023375 |
| Tryptophan | 0.89982 | 0.010347 | 0.02404 |
| Gluconic acid | 1.1258 | 0.013037 | 0.029427 |
| Asparagine | 1.2753 | 0.02567 | 0.05633 |
| Arginine | 0.87285 | 0.034574 | 0.073094 |
| Deoxyadenosine monophosphate | 1.077 | 0.035159 | 0.073094 |

Enrichment pathway analysis of these 35 metabolites revealed that tacrolimus induces changes in the urea cycle, in the amino acid metabolism, in the citric acid cycle and in glutathione metabolism (Figure 2).

### Extracellular metabolomics

Fifty-nine metabolites were detected by LC-MS/MS and, as for intracellular metabolites, heat-map analysis clearly discriminated the control and tacrolimus-treated conditions (Figure 3 and Supplemental table 3). PCA also separated them neatly by means of the first two principal components, 30.3% of the variation being explained by the first and 16.5% by the second (Figure 3).

**Figure 3:**
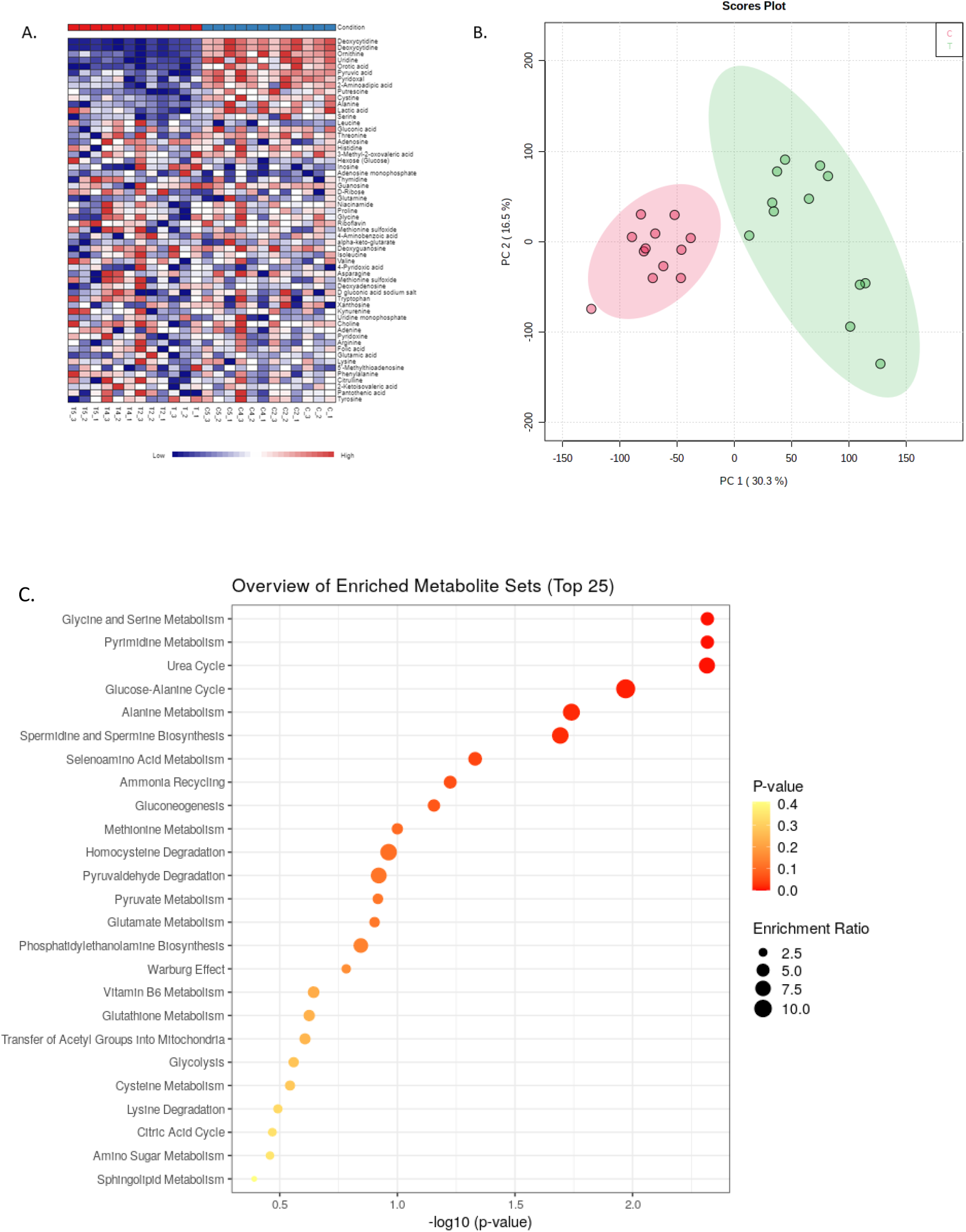
Multivariate exploration of extracellular metabolite variations in response to tacrolimus. Multivariate exploration of extracellular metabolites variations after 24h exposure to tacrolimus (n=12). **A**. Heatmap clustering distinguishing control condition (C) and tacrolimus treated cells (T), based on all metabolites detected by LC-MS/MS analysis. **B**. PCA scores plot showing complete separation between groups (C) and (T) with principal component PC1 and PC2 describing 30.3% and 16.5% of the variations, respectively (0 (red circles): 0.5% ethanol (control); 1 (green circles): tacrolimus 5μM for 24h). **C**. Dot plot of pathway enrichment analysis based on extracellular concentrations of metabolites significantly modified in tacrolimus treated cells as compared to controls (Table 2).

Out of the 59 metabolites detected in the extracellular medium, 17 were significantly impacted (p-value <0.05 and FDR<0.05) by tacrolimus exposure, including amino acids, citric acid cycle intermediates and purine and pyrimidine bases (Figure 3 and Table 2). Enrichment pathway analysis confirmed that the pathways significantly impacted are those of the urea cycle, amino acid metabolism, pyrimidine and glutathione metabolism (Figure 3).

**Table 2.**
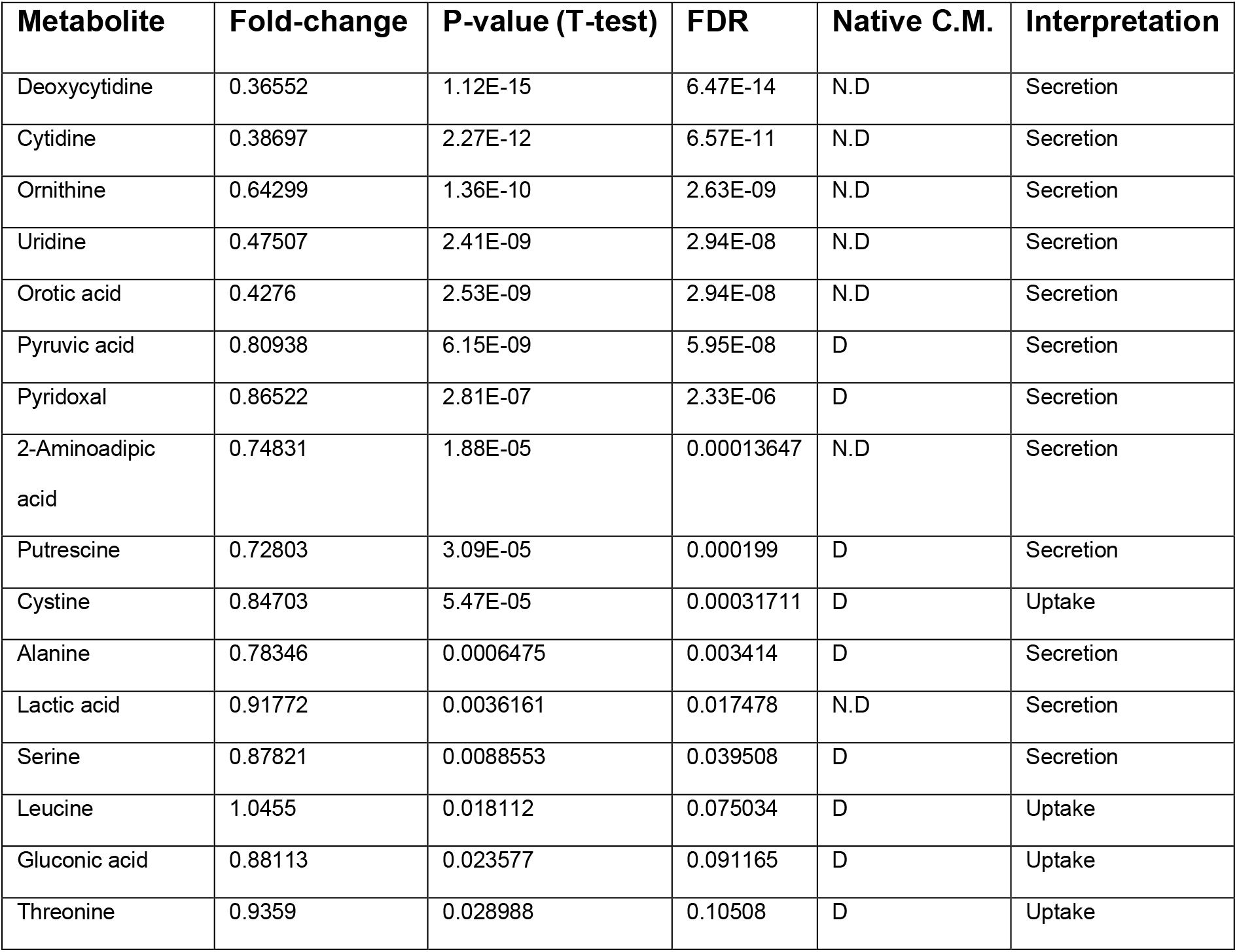
Fold-change of extracellular metabolites significantly influenced by tacrolimus (n=12) (C.M.: culture media, N.D: not detected, D: detected).

| Metabolite | Fold-change | P-value (T-test) | FDR | Native C.M. | Interpretation |
| --- | --- | --- | --- | --- | --- |
| Deoxycytidine | 0.36552 | 1.12E-15 | 6.47E-14 | N.D | Secretion |
| Cytidine | 0.38697 | 2.27E-12 | 6.57E-11 | N.D | Secretion |
| Ornithine | 0.64299 | 1.36E-10 | 2.63E-09 | N.D | Secretion |
| Uridine | 0.47507 | 2.41E-09 | 2.94E-08 | N.D | Secretion |
| Orotic acid | 0.4276 | 2.53E-09 | 2.94E-08 | N.D | Secretion |
| Pyruvic acid | 0.80938 | 6.15E-09 | 5.95E-08 | D | Secretion |
| Pyridoxal | 0.86522 | 2.81E-07 | 2.33E-06 | D | Secretion |
| 2-Aminoadipic acid | 0.74831 | 1.88E-05 | 0.00013647 | N.D | Secretion |
| Putrescine | 0.72803 | 3.09E-05 | 0.000199 | D | Secretion |
| Cystine | 0.84703 | 5.47E-05 | 0.00031711 | D | Uptake |
| Alanine | 0.78346 | 0.0006475 | 0.003414 | D | Secretion |
| Lactic acid | 0.91772 | 0.0036161 | 0.017478 | N.D | Secretion |
| Serine | 0.87821 | 0.0088553 | 0.039508 | D | Secretion |
| Leucine | 1.0455 | 0.018112 | 0.075034 | D | Uptake |
| Gluconic acid | 0.88113 | 0.023577 | 0.091165 | D | Uptake |
| Threonine | 0.9359 | 0.028988 | 0.10508 | D | Uptake |

### Intracellular proteomics

A total of 846 proteins were identified and quantified among the 6 replicates (Supplemental table 4). Only proteins identified using more than 2 peptides were retained. Eleven proteins were found to be differentially expressed and all of them were down-regulated (Figure 4).

**Figure 4:**
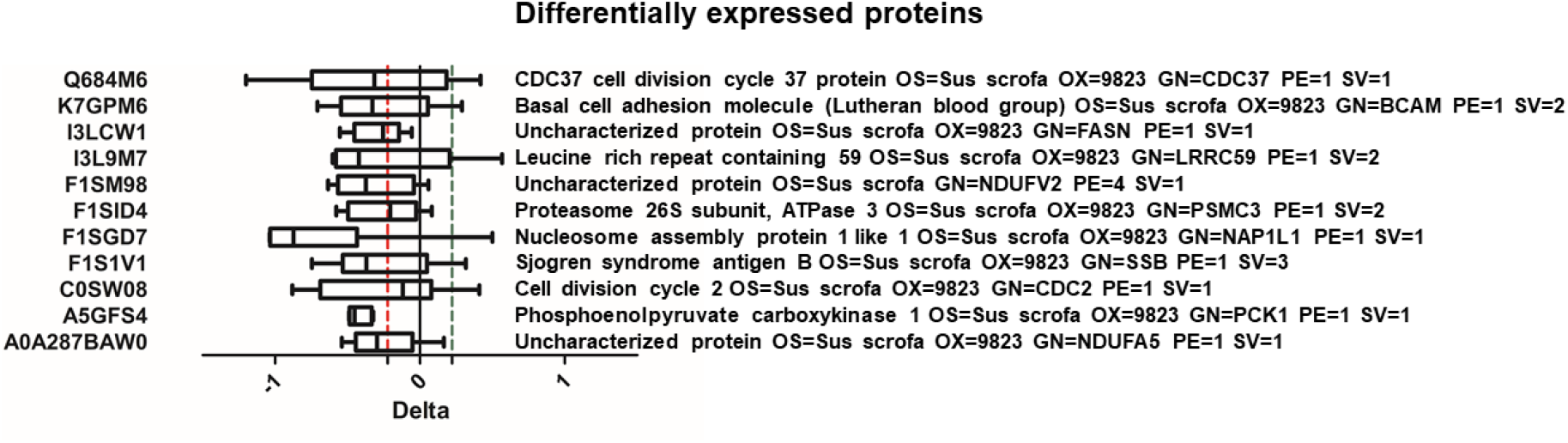
Intracellular variations in proteins induced by tacrolimus. Intracellular concentrations of proteins modified after 24h exposure to 5μM of tacrolimus, as determined by SWATH proteomics analysis (n = 6 independent experiments). Proteins with a mean delta > ln(0.8) or < -ln(0.8) were regarded as differentially expressed.

Two of these proteins are involved in anabolic processes (gluconeogenesis: Phosphoenolpyruvate carboxykinase (PCK-1) “A5GFS4”; Fatty acid synthesis: fatty acid synthase “I3LCW1”), and two others are subunits of the complex 1 involved in the mitochondrial membrane respiratory chain (NADH dehydrogenase [ubiquinone] 1 alpha subcomplex subunit 5 “A0A287BAW0”, NADH dehydrogenase [ubiquinone] flavoprotein 2 “F1SM98”). In our study we will focus on PCK-1 since it was the most down-regulated protein and was consistently down-regulated in the 6 independent experiments, with a mean delta of 0.42 corresponding to a 35% decrease (p< 0.0001). This protein is especially interesting because it can be closely linked to the observed metabolomics perturbations.

### Intracellular targeted transcriptomics

Downregulation of PCK-1 mRNA was confirmed by RT-qPCR (p=0.0329). FBP1, a limiting enzyme in gluconeogenesis was also significantly downregulated (p=0.0031), while the decrease of FBP2, the other limiting enzyme in gluconeogenesis, was not significant (Figure 5.

**Figure 5:**
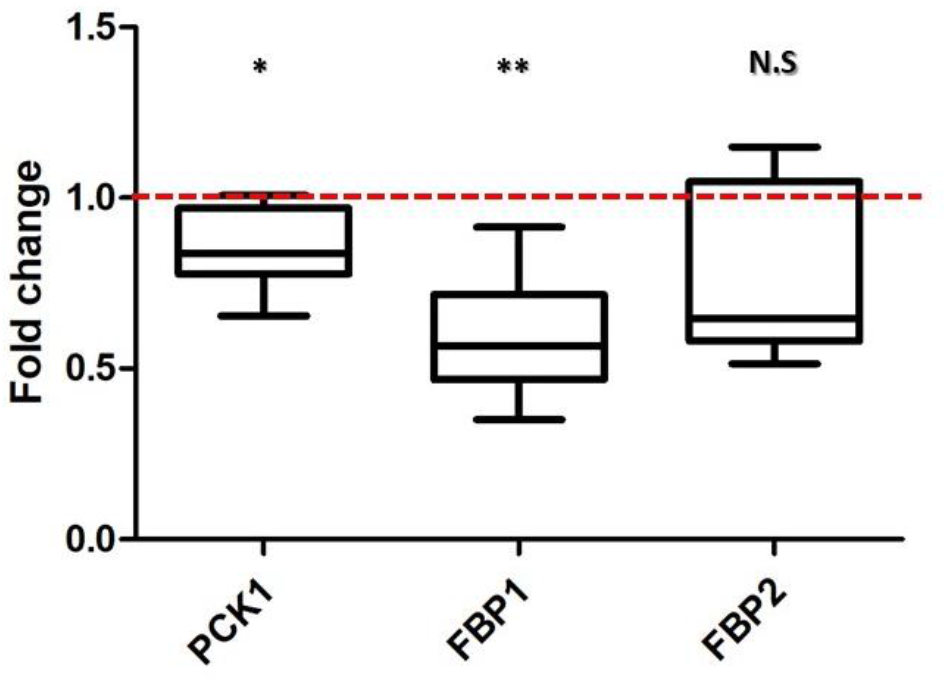
Effect of tacrolimus exposure on the mRNA levels of PCK1, FBP1 and FBP2 in proximal tubular cells. Fold-change of PCK-1, FBP1 and FBP2 mRNAs in cells exposed to 5μM tacrolimus (TAC) with respect to controls (C, exposed to 0.5% ethanol). Targeted gene mRNA signals were normalized to the signal of GAPDH mRNA, as housekeeping gene. * p <0.05; ** p < 0.01 by student *t*-test (n = 6 independent experiments).

## Discussion

This study of the effects of tacrolimus on a porcine cell line (LLC-PK1) suggests that tacrolimus increases the oxidative stress, perturbs the cell energy metabolism and downregulates gluconeogenesis in proximal tubular cells.

As summarized in Figure 6, metabolomics investigations clearly showed increased oxidative stress through a decrease of the total content of intracellular glutathione (*i*.*e*. glutathione, oxidized glutathione and γ-glutamyl-cysteine) in tacrolimus-treated cells. Moreover, the lower extracellular quantity of cystine in the culture medium (Figure 3A, Table 2) suggests an increased uptake of this source of cysteine for glutathione synthesis. Furthermore, increased consumption of folate, known for its free-radical scavenging property, by tacrolimus-treated cells (Table.1) also points towards an oxidative stress. Actually, folate metabolism is a major source of NADPH, a cofactor of glutathione reductase in charge of the regeneration of glutathione from oxidized glutathione (Fan et al., 2014). These results are consistent with previous observations of tacrolimus toxicity mediated by an oxidative stress (Lim et al., 2019a; Zhou et al., 2004) and they validate our experimental setting.

**Figure 6:**
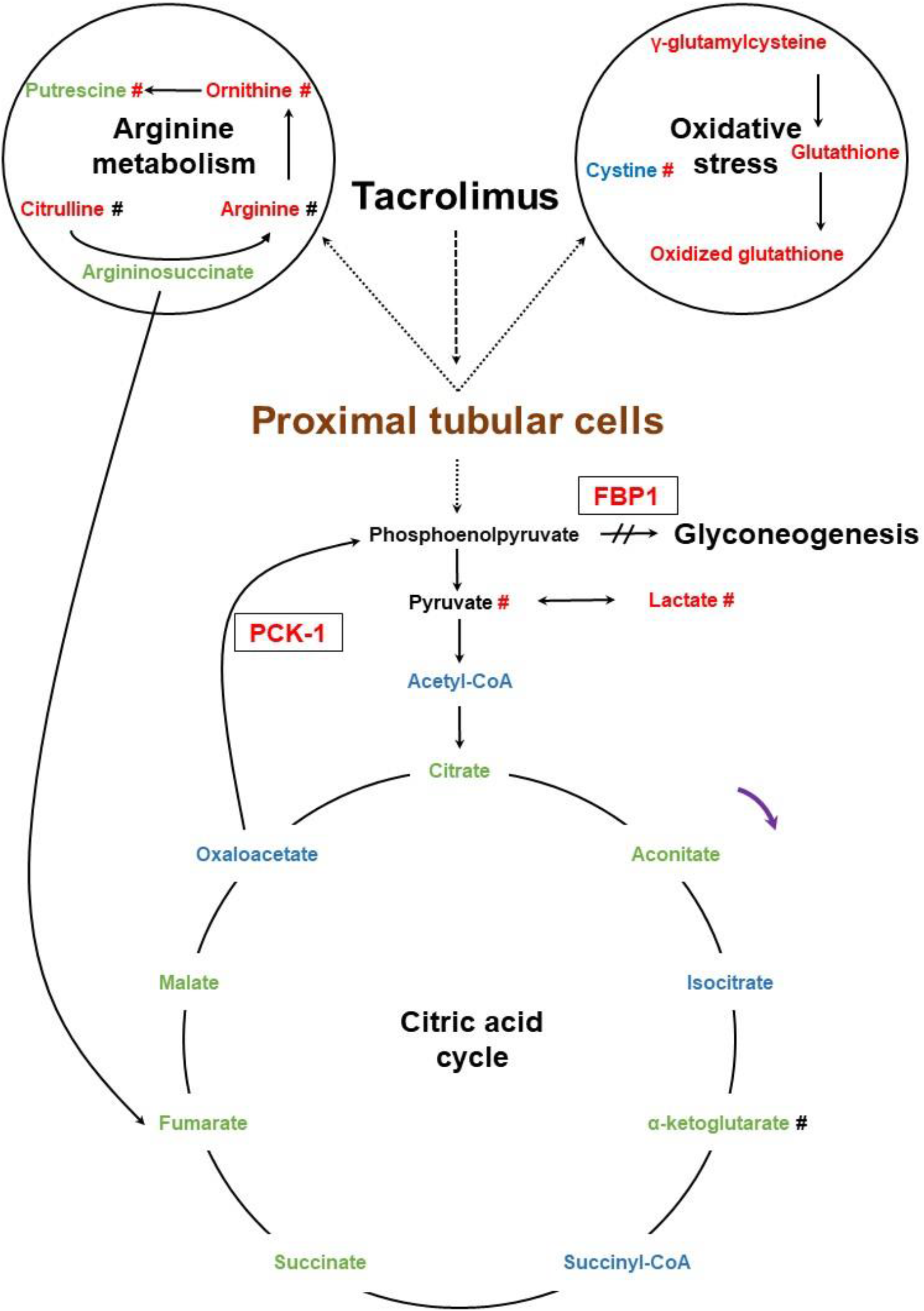
Main pathways modulated by tacrolimus. Intracellular (name) and extracellular (marked with #) metabolites and intracellular proteins (in black boxes) are presented. Decreased metabolites are presented in green, increased metabolites in red, metabolites normally expressed in black and those not detected in blue. Downregulated proteins are presented in red. Purple arrows show cycle directions.

Besides, tacrolimus altered the TCA cycle, as clearly shown by intracellular increase of the TCA cycle intermediates detected (Table 1, Figures 3 and 8). This alteration can be linked with various intracellular events that can induce an accumulation or an increased synthesis of TCA intermediates. Moreover, lactic acid was less excreted and its intracellular concentration decreased (Table 2), which is in favor of an impaired citric acid flux. Our findings suggest possible origins to these TCA cycle perturbations. Similar to folic acid metabolism, the citric acid cycle is a major source of NADPH, which is generated by isocitrate dehydrogenase and malic enzyme and is essential to reduce oxidative stress. Thus, the increase of the citric acid cycle intermediates may be considered as a response to an oxidative stress. Reciprocally, oxidative stress can reduce the activity of some citric acid enzymes like aconitase, oxoglutarate dehydrogenase and succinic dehydrogenase, which may also result in citric acid cycle intermediate accumulation (Tretter and Adam-Vizi, 2005). Our proteomics investigations showed differently expressed proteins, with PCK-1 being the most downregulated (Figure 4). PCK-1 is a limiting enzyme in the gluconeogenesis process that converts oxaloacetate to phosphoenolpyruvate. Interestingly, decreased PCK-1 activity could cause an accumulation of TCA intermediates by limiting oxaloacetate transformation. For instance, PCK-1 -/-mice showed a tenfold increase in malate level in the liver as compared to controls (Hakimi et al., 2005). In addition, this down-regulation of PCK-1 was found in a human hepatic cell line, in primary cultured human β-pancreatic cells and in the liver of mice exposed to tacrolimus (Ling et al., 2020) (Kolic et al., 2020). At the kidney level, a study conducted in rats treated with tacrolimus showed a downregulation of PCK-1 mRNA (Morris et al., 1991). Given the difficulties encountered to find antibodies against porcine proteins, we chose to strengthen our proteomics results by measuring the expression of the corresponding mRNAs. RT-qPCR confirmed that PCK-1 mRNA was down-regulated when cells were exposed to tacrolimus (Figure 5), in line with the above-mentioned studies and our proteomics findings.

TCA cycle impairment can have other origins. One of them is increased glycolysis, essential to produce NADPH that plays an important role in reducing oxidative stress. Another is the stress of the endoplasmic reticulum, known to be implicated in tacrolimus nephrotoxicity and that can increase the citric acid cycle flux mediated by the redox metabolites (Gansemer et al., 2020). Unfortunately, our large, but targeted metabolomics method, may have hampered the identification of key metabolites. Fluxomic approaches could enhance the characterization of tacrolimus-induced TCA cycle perturbations. Monitoring TCA intermediates and glycolysis products during tacrolimus treatment using labelled metabolites could help to determine whether tacrolimus causes an accumulation of TCA intermediates or an increase of the citric acid cycle flux (Lorkiewicz et al., 2019).

Metabolites belonging to the urea cycle were found to be impacted by tacrolimus (Figure 6). Although metabolomics pathway analysis also pointed towards an impact on the urea cycle, proximal tubular cells do not express key urea cycle enzymes, e.g. ornithine transcarbamylase (Levillain et al., 2004). Therefore, tacrolimus disrupted the arginine metabolism. Aspartate, arginosuccinate and fumarate levels were increased in tacrolimus-treated cells, while the levels of arginine, citrulline and ornithine were decreased. On top of the intracellular variations, tacrolimus-treated cells released less ornithine in the medium. In addition, intracellular putrescine, a product of ornithine, increased and its excretion decreased. This may imply a perturbation of the transporter-dependent secretion of putrescine, alone or associated with a potential increase in putrescine synthesis. Interestingly, the accumulation of putrescine can lead to cytotoxicity (Tome et al., 1997; Xie et al., 1997).

In order to investigate if tacrolimus affects only PCK-1 or impacts the whole gluconeogenesis process, mRNA expression of FBP1 and FBP2, two other limiting enzymes in this process, was investigated. Results show that tacrolimus significantly down-regulates FBP1, and probably FBP2 although it did not reach statistical significance (Figure 5). This suggests that tacrolimus affects gluconeogenesis at large and reduces glucose synthesis. A study conducted in mice treated with tacrolimus showed decreased glycemia after 16h of fasting as compared to controls, confirming that tacrolimus affects gluconeogenesis (Ling et al., 2020). This may appear contradictory with the fact that tacrolimus induces diabetes in patients. In fact, tacrolimus diabetogenic effect is probably mostly mediated by its impact on insulin secretion and activity, and not on glucose catabolism (Jindal et al., 1997). All these findings and arguments converge to highlight the effect of tacrolimus on gluconeogenesis in the proximal tubule. Kidneys have gluconeogenesis capacity and play a major role in glucose homeostasis, which may have a systemic impact on patients treated with tacrolimus. Glucose metabolism deregulation induced by tacrolimus in tubular cells, as suggested by down regulation of PCK-1, FBP1 and possibly FBP2, may play a role in tacrolimus nephrotoxicity.

Furthermore, PCK-1 downregulation can impact the acid-base balance. PCK-1 was upregulated in proximal tubular cells in case of acidosis and increased the transformation of oxaloacetate to phosphoenolpyruvate and HCO_3_-(Alleyne and Scullard, 1969; Curthoys and Gstraunthaler, 2001). The latter is released in the blood stream to regulate the acid-base disorder. Tacrolimus favored faster (2 days) systemic acidosis when metabolic acidosis was induced with ammonium chloride in a murine model, whereas acidosis was comparable in mice with or without tacrolimus after chronic exposure to ammonium chloride(7 days) (Mohebbi et al., 2009). This observation was partially explained by alterations of renal acid-base transport proteins. A complementary explanation can now be brought with PCK-1 downregulation by tacrolimus, in as much as PCK-1 is normally upregulated during extracellular and systemic acidosis and *in vitro* hypokalemia induced intracellular acidosis (Alleyne and Scullard, 1969; Curthoys and Gstraunthaler, 2001; Gstraunthaler et al., 2000). Actually, tacrolimus-dependent downregulation of PCK-1 can impact intracellular pH regulation and induce direct toxicity on proximal tubular cells.

All of the metabolomic and proteomic variations described above support tacrolimus-dependent dysregulation of the energy metabolism in tubular proximal cells. These cells, due to their functions involving a large number of energy-dependent transports, can be largely impacted by these perturbations, which could be root mechanisms in the tubular toxicity of tacrolimus. In addition, PCK-1 downregulation can have an impact beyond the energy metabolism and may affect specific tubular functions such as glycemia regulation and acid-base balance.

It is worth mentioning that the tacrolimus concentration used in this study was much higher than trough tacrolimus whole blood concentrations found in treated patients (typically 4-15 ng/ml). However, the concentration used (5μM), did not show any cytotoxic effect after 24h incubation and it is at the lower end of the concentrations previously used to investigate tacrolimus nephrotoxicity *in vitro*, which can be up to 74 μM (Bennett et al., 2016; González-Guerrero et al., 2013; Lim et al., 2019a, 2019b, 2017; Yu et al., 2019; Zhou et al., 2004). Still, toxicity findings made at these relatively high concentrations could be validated in animal models of chronic tacrolimus nephrotoxicity (Lim et al., 2019b; Yu et al., 2019). This suggests that our results, obtained at intermediate concentrations, may raise pertinent hypotheses regarding tacrolimus toxicity pathways. Because tacrolimus nephrotoxicity (apart from its vasoconstrictive properties) is a delayed and chronic process, its *in vitro* exploration would ideally require a very long incubation time, but this is hardly feasible. There is no example in the literature we know of that employed a different strategy than using high concentrations of tacrolimus for short time-periods. Finally, the variations reported here were observed at or after 24h incubation and a kinetics study might allow better understanding of tacrolimus underlying mechanisms.

## Conclusion

This multi-omics study conducted on a proximal tubular cell line shows that acute tacrolimus induces alterations in the energy and glucose metabolisms and increased oxidative stress, which can impact glycemia regulation and disrupt the intra-extracellular acid-base balance. These short-term modifications might be implicated in tacrolimus chronic nephrotoxicity.

## Supporting information

supplemental data Table 1. RT-qPCR primers

supplemental data table 2. intracellular metabolomics data

supplemental data table 3. extracellular metabolomics data

supplemental data table 4.Proteomics data

## Abbreviations

PCK-1: phosphoenolpyruvate carboxykinase 1
FBP1: Fructose biphosphatase 1
FBP2: Fructose biphosphatase 2
LLC-PK1: Lilly Laboratories Cells Porcine Kidney 1
LC-MS/MS: liquid chromatography coupled to tandem mass spectrometry
MTS: 3-(4,5-dimethylthiazol-2-yl)-5-(3-carboxymethoxyphenyl)-2-(4-sulfophenyl)-2H-tetrazolium
NADPH: Nicotinamide adenine dinucleotide phosphate
PCA: Principal component analysis

## Acknowledgments

The authors thank Jean-Sebastien Bernard for helping with the experiments.

## Author contribution

H.AD, Q.F, P.M, M.E participated in research design. H.AD conducted the experiments. H.AD, FL.S, E.P, H.A, contributed to the experiments. H.AD, Q.F, CC.B performed data analysis. H.AD, P.M, ME, FL.S, E.P, Q.F CC.B, H.A wrote or contributed to the writing of the manuscript.

## Footnotes

Reprint requests should be sent to Prof. Pierre Marquet, INSERM UMR1248 “Pharmacology & Transplantation”, Faculty of Medicine, 2 rue du Dr. Marcland, 87042 Limoges, France.

