## supplemental data Table 1. RT-qPCR primers for "Multi-omics investigation of tacrolimus nephrotoxicity"

Table 1: Primers sequences used for RT-qPCR

| Gene | Sequence 5’=>3’ | |
| --- | --- | --- |
| PCK-1 | Forward | TGGCCATGATCCAAAGACCG |
|  | Backward | GGCAGAACCTCCCAGCTTTA |
| FBP1 | Forward | ACGGTGCTACCTTATTCTGGC |
|  | Backward | GGTAGCACACGGCATTCACA |
| FBP2 | Forward | GCAGGCAGGTAGTGAGTCAG |
|  | Backward | GACTTGCCTCCTGCTTCTCA |
